# Single-cell multiomics reveals the oscillatory dynamics of mRNA metabolism and chromatin accessibility during the cell cycle

**DOI:** 10.1101/2024.01.11.575159

**Authors:** Maulik K. Nariya, David Santiago-Algarra, Olivier Tassy, Marie Cerciat, Tao Ye, Andrea Riba, Nacho Molina

## Abstract

The cell cycle is a tightly regulated process that requires precise temporal expression of hundreds of cell cycledependent genes. However, the genome-wide dynamics of mRNA metabolism throughout the cell cycle remain uncharacterized. Here, we combined single-cell multiome sequencing, biophysical modeling, and deep learning to quantify rates of mRNA transcription, splicing, nuclear export, and degradation. Our approach revealed that both transcriptional and post-transcriptional processes exhibit distinct oscillatory waves at specific cell cycle phases, with post-transcriptional regulation playing a prominent role in shaping mRNA accumulation. We also observed dynamic changes in chromatin accessibility and transcription factor binding footprints, identifying key regulators underlying the oscillatory dynamics of mRNA. Taken together, our approach uncovered a high-resolution map of RNA metabolism dynamics and chromatin accessibility, offering new insights into the temporal control of gene expression in proliferating cells.

**Highlights:**

- FourierCycle combines single-cell multiome sequencing, deep learning, and biophysical modeling to quantify gene-specific rates of mRNA metabolism during the cell cycle
- Rates of mRNA transcription, nuclear export, and degradation show gene-specific oscillatory waves at distinct cell cycle phases.
- Post-transcriptional regulation, including mRNA degradation and nuclear export, plays a prominent role in shaping mRNA accumulation during the cell cycle
- Dynamics of chromatin accessibility and transcription factor binding footprints uncover key regulators underlying the transcriptional control of gene expression

**Graphical abstract:** 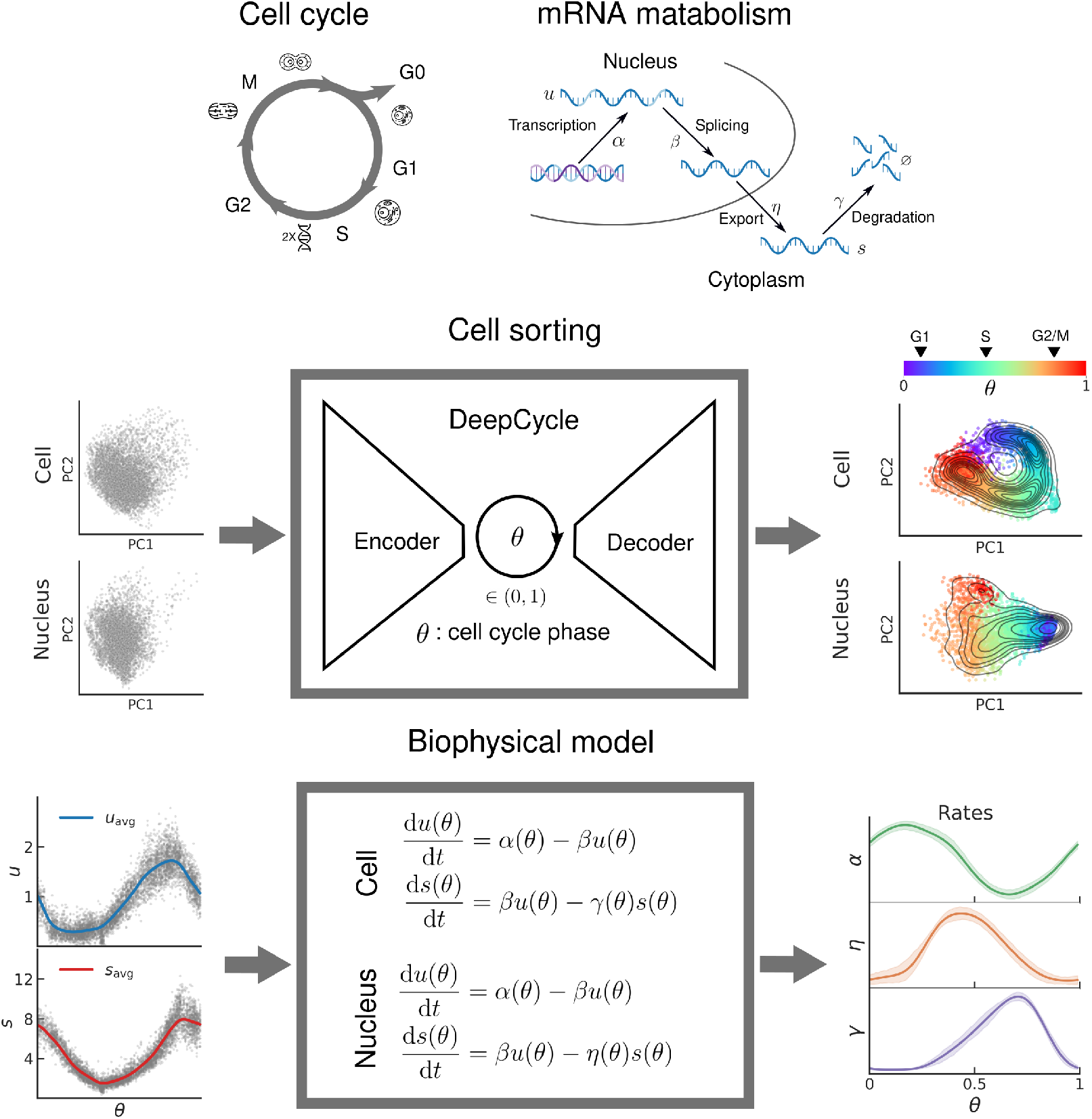

## Introduction

The cell cycle is a highly regulated process that ensures the accurate replication and transmission of genetic information from one generation of cells to the next. It is a fundamental biological process and plays a crucial role in development, growth, and maintenance of all living organisms, and its dysregulation can lead to a number of pathologies such as autoimmune diseases, neurodegenerative diseases, and cancer [1–8]. In general, cell cycle proceeds through four phases, G1, a period when cell duplicates essential cellular components, S, a period of DNA duplication, G2, when organelles and proteins for cell division develop, and M phase that completes the process [1]. Unimpeded progression through the cell cycle requires precise and regulated deployment of hundreds of proteins, and thus represent one of the most complex and tightly spatiotemporally controlled process in biology [1–8]. The levels of these proteins are in part controlled by periodic expression of their mRNAs during the cell cycle, often observed as waves of gene expression [9–11]. However, studies of the mRNAs and how they emerge, change and dissipate during different phases of cell cycle at high temporal resolution has remained technically challenging.

Recently, RNA velocity, *i*.*e*. the time derivative of the gene expression state, was introduced as a framework to obtain the systems-level dynamics of gene expression from static snapshots of transcriptional states measured at single cell resolution [12–15]. RNA velocity depends on the four main steps in mRNA metabolism: (1) transcription, which generates pre-mRNA (unspliced mRNA); (2) pre-mRNA splicing to obtain mature mRNA (spliced mRNA); (3) the mature mRNA export into the cytoplasm for protein synthesis; and (4) mRNA degradation. In this context, measuring unspliced and spliced mRNA levels in cells at single-cell resolution can provide a dynamical picture of cellular processes [12–15]. However, this approach does not provide a way to gain insights into the dynamics of the cell cycle. Furthermore, although computational tools such as Cyclum and Revelio have been developed to infer the cell cycle phase based on scRNA-seq data, these methods do not take into account RNA velocity and offers no insight into RNA metabolism [10, 11]. To begin to address these limitations, we recently developed DeepCycle, a deep learning tool that uses single-cell RNA sequencing (scRNA-seq) to map the gene expression profiles to a continuous latent variable, *θ*, representing the cell cycle phase [16]. We used DeepCycle to study dynamics of gene expression at high temporal resolution in embryonic and somatic cells, revealing major waves of RNA accumulation during the cell cycle and identifying key transcriptional regulators involved in cell cycle progression. This led to the question: how are the processes that govern RNA accumulation, such as transcription, nuclear export, splicing, and degradation, temporally regulated during the cell cycle on a genome-wide scale? Here, we address this by combining single-cell RNA sequencing, single-nucleus multiome sequencing, DeepCycle, and biophysical model construction. Our model, which we call FourierCycle, predicts that cell cycle depends on transcription, nuclear export, and degradation rates for every gene, revealing waves of transcriptional and post-transcriptional regulation during the cell cycle. Furthermore, our method allowed us to investigate changes in chromatin accessibility at the gene level and their potential connection to transcription. Remarkably, we identified key transcription factors (TFs) and the dynamics of their binding activities during the cell cycle by examining changes in chromatin accessibility footprints near their binding sites. In summary, we developed a framework that combines state-of-the-art sequencing technologies with deep learning and biophysical modeling to uncover a global picture of transcriptional and post-transcriptional regulation of mRNA accumulation and chromatin accessibility dynamics throughout the cell cycle.

## Results

### DeepCycle sorts single cells and single nuclei according to their cell cycle progression

We cultured mouse embryonic stem cells (mESCs) in leukemia inhibitory factor (LIF), supplemented with an inhibitor of the mitogen-activated protein kinase enzyme MEK, and a glycogen synthase kinase-3 (GSK3) inhibitor (details provided in Methods). This treatment promotes the maintenance of a self-renewal cell state, reducing heterogeneous expression of pluripotency factors and spontaneous differentiation, thereby generating an asynchronous population of proliferating cells [17, 18]. We previously performed scRNA-seq to obtain the transcriptomes of whole mESCs [11]. To complement this dataset with nuclear RNA expression and chromatin accessibility information, we employed the multiome protocol that involved simultaneously performing single-nucleus RNA sequencing (snRNA-seq) and singlenucleus Assay for Transposase-Accessible Chromatin using sequencing (snATAC-seq) on the same cells [19]. We profiled around 5,000 cells in each experiment at high sequencing depth, revealing the gene expression of roughly 12,000 genes. To obtain an overall view and comparison between the single-cell and single-nucleus data, we calculated the average reads of unspliced and spliced RNA across genes and across cells (or nuclei) (**Figure 1A – 1D**). Comparing the unspliced and spliced reads in the cell with ones in the nucleus revealed strong correlation between the mRNA reads obtained from the nuclear data and cell data (*r* = 0.86 for unspliced and 0.78 for spliced), suggesting a high degree of coherence in the measurements (see **Figure S1** for corresponding figures for replicate 2). The points in the scatterplot in **Figure 1C** are equally distributed across the *x* = *y* line, suggesting that most of the unspliced molecules of the cell reside in the nucleus, whereas most of the points in the scatterplot in 2D are below the *x* = *y* line, suggesting that most of the spliced molecules are present in the cytoplasm of cell, in agreement with what we expected to see.

**Figure 1.**
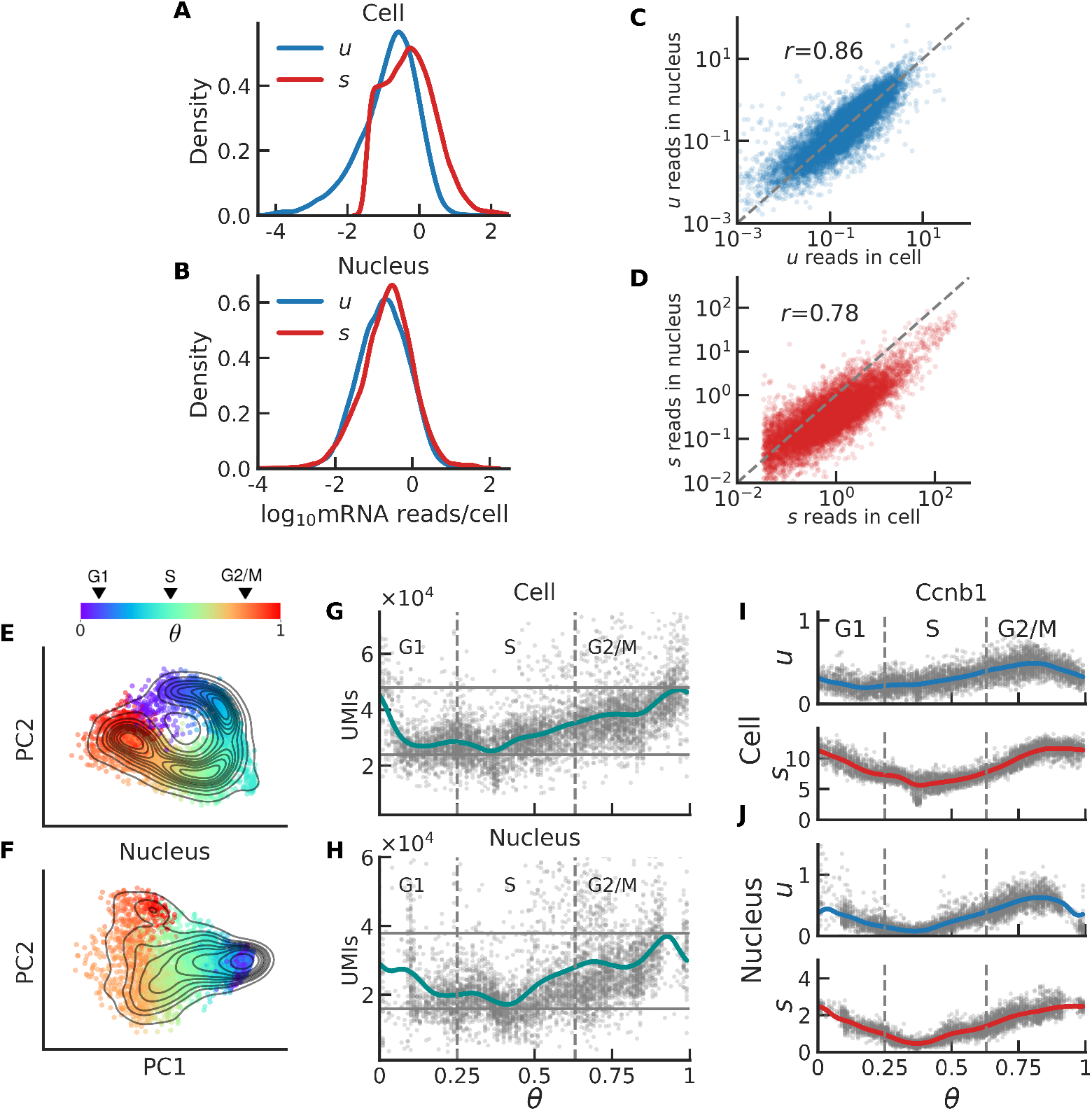
DeepCycle infers the cell cycle phase of both single cells and single nuclei. **A** and **B**: Distribution of unspliced (blue) and spliced (red) reads per cell, across all genes in the cell and the nucleus respectively. **C** and **D**: Correlation between the reads in the cell versus those in the nucleus for the unspliced (blue) and spliced (red) molecules respectively, every dot represents a gene. **E** and **F**: Visualization of the cycling dynamics based on the sequencing profiles of the cells and isolated nuclei respectively, every dot represents a cell, the color indicates the progression through the cell cycle–early G1 (in purple) through mitotic (in red), **G** and **H**: Unique molecular identifiers (UMIs), a proxy for the total mRNA content in the cell and the nucleus respectively, against the cell cycle phase *θ*; there is a two-fold change in the total mRNA content, both in the nucleus as well as in the cell, during the cell cycle. **I** and **J**: Expression dynamics of the unspliced and spliced reads for Ccnb1 in the cell and the nucleus respectively. The cell cycle phase transitions are represented with grey dotted lines and obtained using our previously described approach [11]. The grey dots get scarce near *θ* approaches 0 or 1, this is likely because the nuclear envelope breaks down during mitosis and the sequencing technique is unable to profile these nuclei.

Next, we used DeepCycle and both the scRNA-seq and snRNA-seq data from mESCs and determined the cell cycle phase of each cell and nucleus. Our approach effectively revealed the cell cycle manifold embedded in gene expression space, as depicted in the projection onto the two main principal components (PC) for whole cells and for the nucleus (**Figure 1E – 1F**). We observed that for the single cell data, the cells are accurately positioned along the dynamic cycling trajectory and distinguishable based on their progression through the cell cycle (**Figure 1E**). For the single nuclei, although the cell cycle manifold may not appear to be cyclic, the nuclei cluster well according to their progression through the cell cycle in the correct order (**Figure 1F**). It is worth noting that the cells approaching mitosis and the cells in early G1 phase were filtered out as they often appear as extreme outliers in the PC-space (see **Figure S1**). The number of unique molecular identifiers (UMIs) [20, 21], which can be interpreted as total mRNA content of the cell, show a two-fold change as the cells progress through the estimated cell cycle phase (**Figure 1G**). This is the expected behavior, assuming that the concentration of mRNA remains constant during the cell cycle [22]. Importantly, the total mRNA content within the nucleus also exhibits a similar two-fold increase as observed in the whole cell during the cell cycle (**Figure 1H**). Lastly, expression dynamics of the unspliced and spliced reads for Cyclin B1 (Ccnb1), a crucial regulator of the G2/M transition during the cell cycle [23, 24], showed gene-specific dynamics of unspliced and spliced levels of mRNA in the cell and the nucleus (**Figure 1I – 1J**). We observed that the unspliced and spliced RNA nuclear levels of Ccnb1 mirror the dynamics observed in the entire cell.

In summary, we obtained a unique scRNA-seq and snRNA-seq datasets at a high sequencing depth and showed that DeepCycle accurately assigns a continuous phase to both single cells and single nuclei, indicating their progression through the cell cycle.

### Kinetic modeling reveals cell cycle-dependent mRNA transcription and degradation rates

DeepCycle reveals oscillatory patterns in mRNA accumulation throughout the cell cycle, prompting an investigation into the underlying processes of mRNA metabolism responsible for these dynamics. To address this question, we developed FourierCycle, a biophysical model of mRNA metabolism. As mentioned earlier, the key steps involved in mRNA metabolism include transcription, splicing, nuclear export, and degradation. These processes were modeled using a system of coupled differential equations with periodic functions to capture cell cycle-dependent rates. We derived a pseudo-analytical solution using Fourier series expansion (details provided in the supporting information). Finally, we used normalized unspliced and spliced reads, along with the cell cycle phase as observable variables, to fit the fundamental rates of RNA metabolism (details in the Methods section).

Our biophysical model predicted the dynamics of the unspliced and spliced mRNA levels as well as the dynamics of the transcription and degradation rates in good agreement with experimental data (see representative genes Ccne1 and Wee1; **Figures 2A** and **2B**). Ccne2 encodes the G1/S-specific cyclin-E2 protein, essential for cell cycle control during the late-G1/early-S phase [25, 26], whereas Wee1 acts as a negative regulator of entry into mitosis [27, 28]. In agreement with these established functions, we observed a significant increase in the Ccne2 transcription rate during the G1/S transition (**Figure 2A**). Additionally, we noted that the degradation rate remains relatively constant, suggesting that the mRNA level of Ccne2 is primarily regulated by transcription. Also in agreement with the literature, Wee1 showed a sharp increase in mRNA degradation during the M/G1 transition, while its transcription rate did not fluctuate much relative to the baseline (**Figure 2B**). This suggests that, in the case of Wee1, mRNA expression is mainly regulated post-transcriptionally.

**Figure 2.**
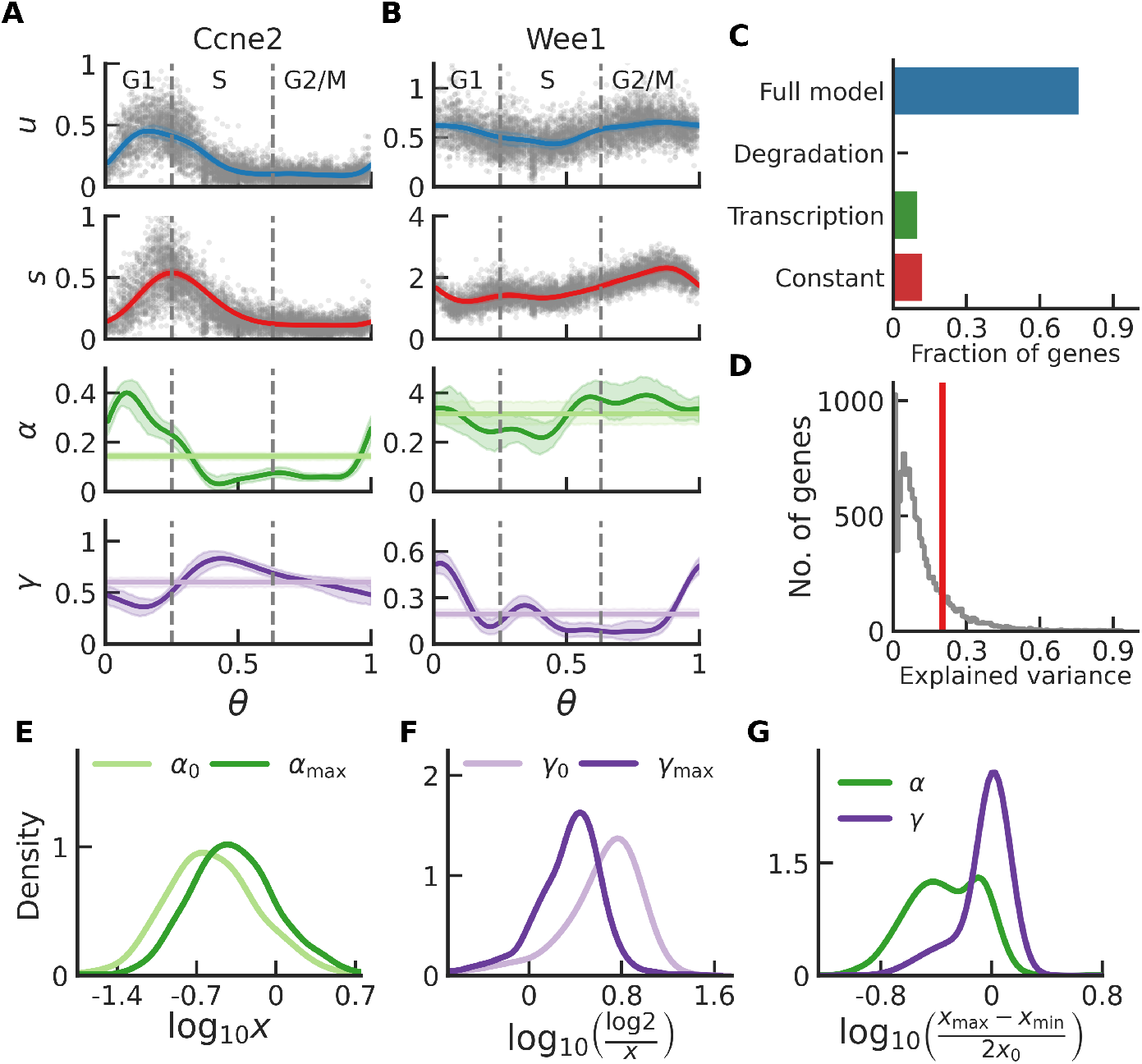
Genome-wide estimation of global transcription and degradation rates and characterization of transcriptional and post-transcriptional regulation. **A** and **B** (top to bottom): number of unspliced reads (*u*), number of spliced reads (*s*), transcription rate (*α*), degradation rate (*γ*) as function of the cell cycle phase *θ*. Grey dots represent single cells, solid lines show model predictions, and shaded regions are 95% confidence intervals from bootstrapping. **C:** Barplot of fraction of genes selecting alternative models based on the Bayesian Information Criterion–full model (blue), model with cell cycle dependent degradation and constant splicing and transcription (purple), model with cell cycle dependent transcription and constant splicing and degradation (green), model with constant transcription, splicing, and degradation (red), for the vast majority of the genes the full model is preferred. **D:** Histogram of the explained variance in spliced counts across all the genes; red line is at *x* = 0.2, the threshold set for filtering genes. **E:** Distribution of transcription rate, in arbitrary units, across filtered genes, *α*_0_ denotes baseline transcription (light green) whereas *α*_max_ denotes maximum transcription rate (green), F: Distribution of degradation half lives, in hours, across filtered genes, *γ*_0_ denotes baseline degradation rate (light purple) whereas *γ*_max_ denotes maximum degradation rate. **G:** Fold change in amplitude of rates, (*x*_max™_ *x*_min_)*/*2, over the baseline rate for transcription (in green) and degradation (in purple). For most of the genes, the changes in degradation rates are more prominent than those in transcription rates compared to the corresponding baseline rates.

Due to the high noise level of single-cell sequencing technologies, many genes do not fit the model with the desired level of statistical significance. Thus, we used the explained variance, *r*^2^, as a measure of the discrepancy between the model and the data. By filtering genes with *r*^2^ *>* 0.2, approximately 1,800 out of the 12,000 profiled genes met this threshold (**Figure 2D**). All downstream analyses were restricted to these genes. To determine gene-specific preferences for transcriptional or post-transcriptional regulation genome-wide, we constructed three alternative models: one where only transcription is cell cycle-dependent (splicing and degradation remaining constant); another where only degradation is cell cycle-dependent (transcription and splicing remaining constant); and a third where transcription, splicing, and degradation all remain constant during the cell cycle. We fitted these models to the sequencing data and used Bayesian information criterion (BIC) to perform model selection for every gene. We observed that the behavior of 67.5% of the genes was captured by a full model with cell cycle-dependent transcription as well as degradation; 19.2% of genes agree with the model where only transcription is cell cycle dependent; 12.5% of the genes prefer the model with no cell cycle dependence; and only 1% of the genes prefer the model where only degradation is cell cycle dependent (**Figure 2C**). Overall, we were able to fit our biophysical model of mRNA metabolism to thousands of genes.

We investigated the genome-wide synthesis and degradation rates for all the genes satisfying the explained variance cutoff mentioned above. We observed that the baseline transcription rates and degradation rates in mESCs span a large dynamic range across genes–more than two orders of magnitude (**Figure 2E** and **2F**). Since the model was fitted to relative quantifications of unspliced and spliced reads, the numerical values of transcription rates in **Figure 2E** do not reflect the absolute number of molecules synthesized per hour; however, fold changes with respect to the baseline provide information on the transcriptional resources dedicated to a particular gene during cell cycle progression. On the contrary, the degradation half-lives were expressed in units of hours, assuming a typical cell cycle duration of 14 hours (**Figure 2F**) [11]. We found that the mean of the baseline degradation half-lives was around 4.5 hours, whereas the mean of the maximum degradation half-lives was 2.7 hours. These values overall agree with previously reported estimates of mRNA degradation rates [29, 30].

We also calculated the difference in time between peak spliced and peak unspliced levels. For the majority of the genes, the time difference is less than one-eight of a cycle, which translates to less than 1.75 hours (**Figure S2**). To gain insight into the fold changes in the rate parameters, we calculated the amplitude of the rates, *i*.*e*. (max − min)/2, and compared it to the baseline rates (**Figure 2G**). For most of the genes, the fold-change in amplitude for mRNA degradation is higher than transcription. Model selection using BIC suggests that regulation of mRNA metabolism requires both transcription and degradation to be cell cycle dependent; however, for the majority of genes, the changes in mRNA degradation rates contribute more to regulating gene expression dynamics than the changes in transcription. Therefore, mRNA post-transcriptional regulation has a prominent role in shaping mRNA accumulation during the cell cycle.

### Waves of transcriptional and post-transcriptional regulation during the cell cycle

After investigating the global trends in transcription and degradation rates, we examined the timing of peak rates of transcription and degradation during the cell cycle for the filtered genes. We observed distinct waves of mRNA synthesis during the cell cycle (**Figure 3A**), with 762 genes (44.3% of the 1778 filtered) achieving peak transcription during the G1 phase, 481 genes (28.0%) during the S phase, and 477 genes (27.7%) during the G2/M phase. In terms of mRNA degradation, we found that 820 (47.7%) genes achieved peak degradation during the G1 phase, 707 (41.1%) genes during the S phase, and 193 (11.2%) genes during the G2/M phase (**Figure 3B**). Interestingly, we observed a prominent wave of maximum transcription in the early G1 phase (**Figure 3A**), and waves of degradation concentrated mainly during G1 and S phases (**Figure 3B**). Next, we normalized the predicted transcription and degradation rates and arranged genes based on the timing of their peak transcription (**Figure 3C** and **3D**), observing the same wave patterns. However, each of these waves consists of a distinct set of genes. For example, during the G1, the mESCs upregulate genes involved in fundamental processes essential for cell growth and proliferation (*e*.*g*. amine metabolism, DNA replication initiation, and cell cycle G1/S phase transition (**Figure 3E**). On the other hand, S phase involves peak transcription of genes involved in preparing cells for mitosis (*e*.*g*. nucleosome organization, regulation of cell cycle G2/M transition, establishment of spindle orientation, **Figure 3E**), and G2/M phase is associated with synthesis of genes involved in a variety of mitosis related pathways (**Figure 3E**). The enrichment analysis of the degradation waves revealed many of the same processes; however, they tended to peak at different times consistent with the need to remove transcripts of proteins that are no longer functionally needed (**Figure 3F**). As such, the degradation wave during the G1 phase consists of genes that are involved in mitotic regulation, as genes produced during the G2/M phase are degraded upon entering the G1 phase, when they are no longer needed. Similarly, during the S phase, the mECSs degrade genes involved in processes such as DNA replication initiation and G1/S phase transition, which achieved peak transcription during the G1 phase. Overall, these results suggest that waves of transcription and mRNA degradations occur at well-defined points of the cell cycle. We speculate that they enable the cell to produce transcripts (ultimately proteins) needed at specific points of the cell cycle *en masse*, and put a stop to their activity by waves of mRNA degradation.

**Figure 3.**
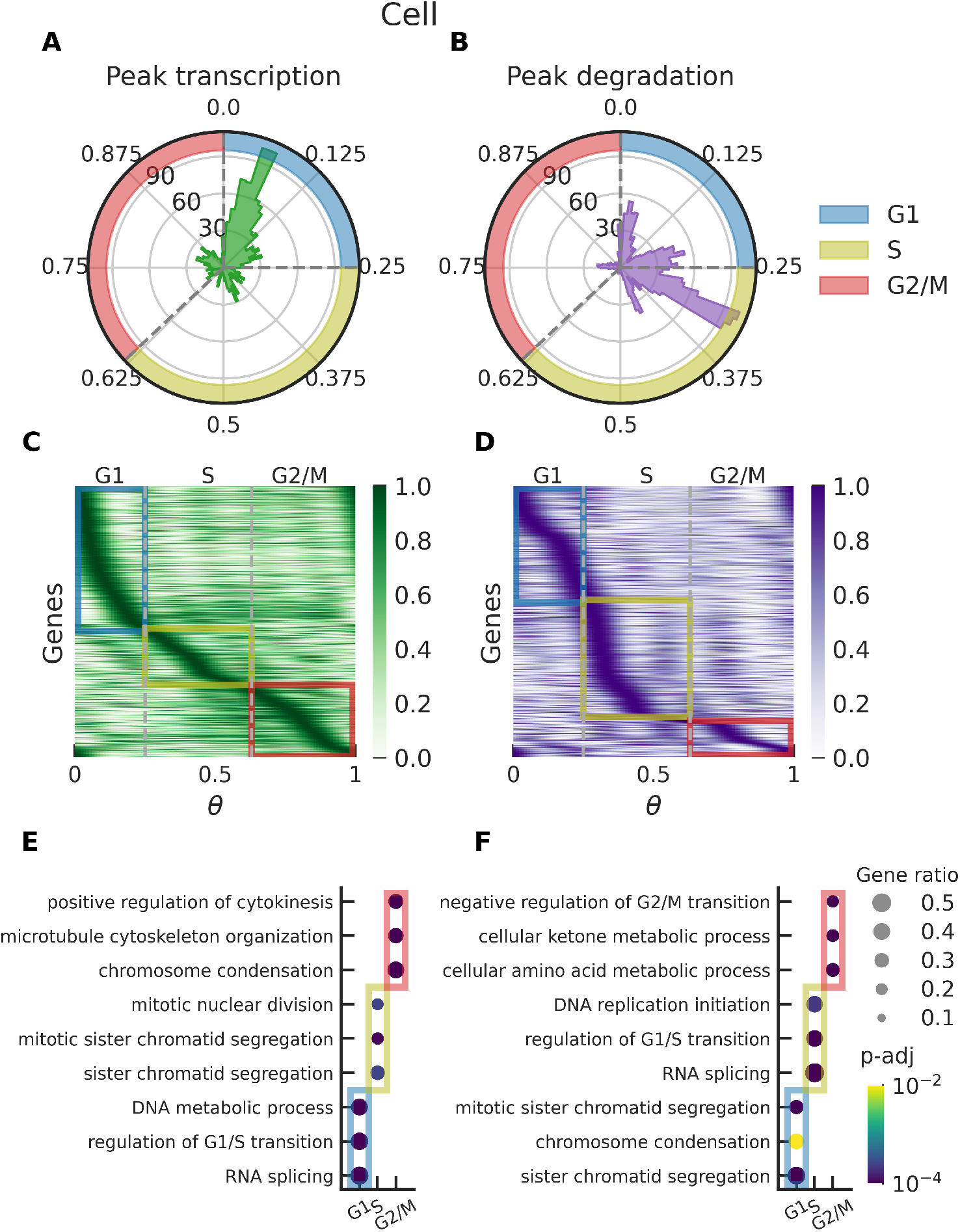
Waves of transcription and degradation and the biological processes involved during the cell cycle. **A** and **B:** Polar histograms of the cell cycle phase at which a gene achieves the highest transcription rate and the highest degradation rate, respectively. **C** and **D:** Heatmaps of transcription and degradation, respectively, during the cell cycle. The rates are normalized using max-min normalization, and the genes are arranged based on the timing of the maximum rate. Genes that achieve their peak transcription and degradation rates during G1, S and G2/M phases are highlighted in light blue, olive, and pink, respectively. **E** and **F:** Gene set enrichment analysis [31, 32] for genes that achieve peak transcription and degradation respectively; gene sets were compared against GO biological processes 2018 (in mouse) size of the dot represents the extent of overlap between the gene set and the GO terms, the color represents the adjusted *p*-value.

### Genome-wide estimates of mRNA nuclear export rates and waves of nuclear export

Nuclear mRNA export is another process that could affect mRNA metabolism. To measure mRNA export rates and their changes during the cell cycle, we constructed a biophysical model that attributes changes in abundance of mRNA in the nucleus to transcription, splicing, and nuclear export, denoted by *η*. This model makes an assumption that nuclear export exerts much stronger influence over mRNA abundance in the nucleus than degradation, which may be a reasonable scenario given that nuclear export is a faster process than nuclear degradation [33]. As seen in the case of transcription and degradation, nuclear retention halflives span a large dynamic range, with the mean baseline half-life of 1.5 hours and the mean maximum half-life of 36 mins (**Figure 4A** and **4B**). This is in agreement with previously reported values [33–35]. The distribution of foldchange in amplitude over the baseline for nuclear export half-lives was between 0.001 to 4.11 (**Figure 4B**), indicating that for some genes, the change in export rate can play a significant role in the dynamics of mRNA accumulation during the cell cycle. We found that majority of genes (67.6%) prefer the full model (transcription, splicing and nuclear export) (**Figure 4C**), indicating that nuclear mRNA retention may be regulated during the cell cycle. For 19.0% of the genes, we found that only transcription was cell cycle-dependent, and 12.5% of the genes preferred a model with constant rates. We also calculated the explained variance and filtered genes with *r*^2^ *<* 0.2, with roughly 2,000 (out of 12,000) satisfying the criterion (Figure S3F and S3I). This number is higher than what we observed in our model that considers degradation (1,800 genes out of 12,000; see above). This could be explained by the superior capture efficiency of intronic reads (unspliced mRNA) in the case of the snRNA-seq when compared to scRNA-seq, which directly impacts the RNA-velocity framework. We also examined the timing of peak transcription and export rates. Strikingly, our method uncovers very prominent waves of nuclear export during the cell cycle (Figure 4D and Figure S5). Of the 2170 genes that satisfied the explained variance criterion, 805 (21.4%) genes achieved peak export during the G1 phase, 1351 (35.9%) genes during the early-S phase, and 1608 (42.7%) genes during the G2/M phase (**Figure 4D** and **4E**). Taken together, we identified evidence that mRNA nuclear export was cell cycle dependent. We also observed waves of maximal rates of nuclear export at different points during the cell cycle, most notably the early-S phase and the late G2/M phase.

**Figure 4.**
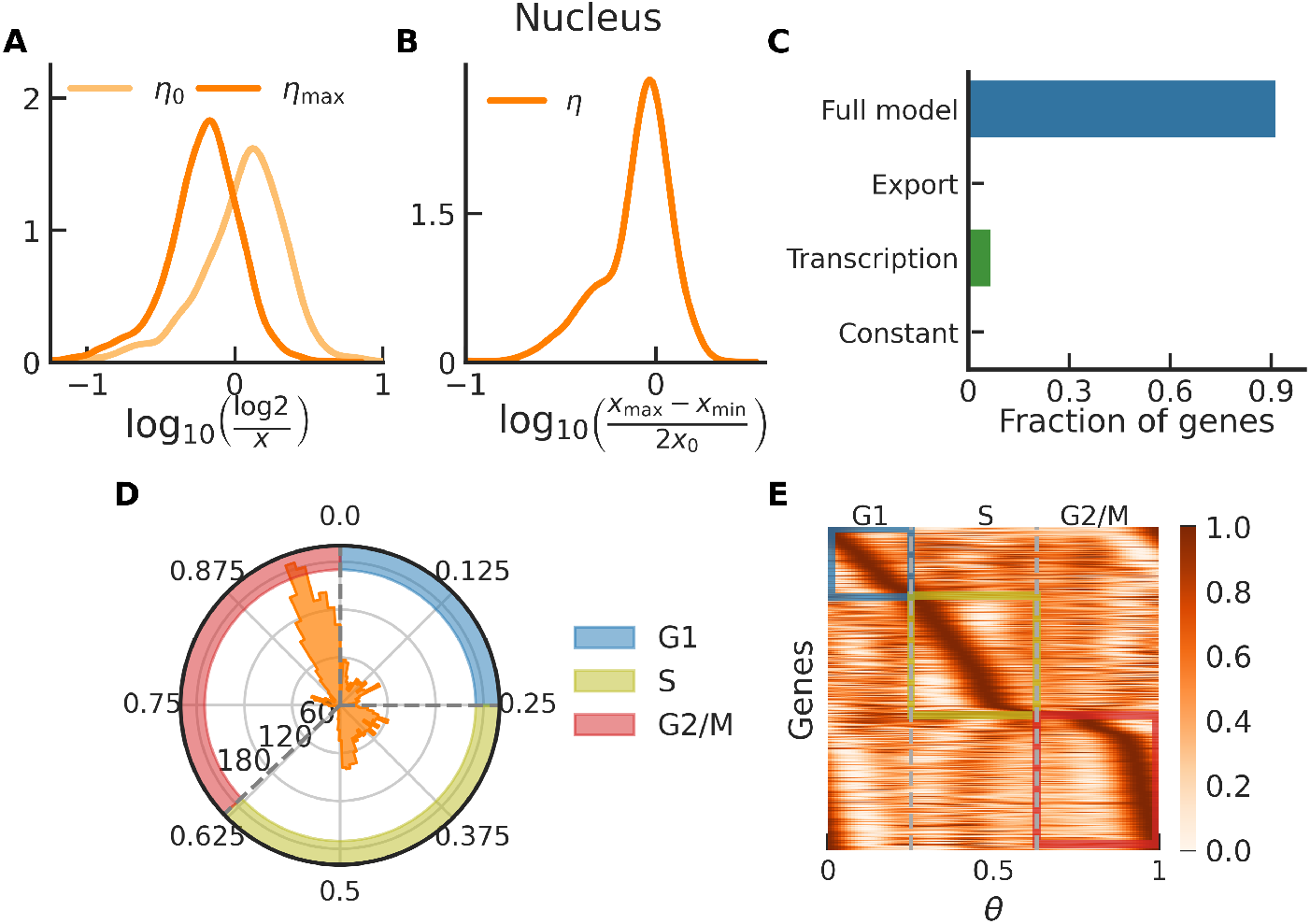
Genome-wide estimation of global export rates and characterization of waves of export during the cell cycle. **A:** Distribution of export half-lives, in per hour, for baseline export rate (light orange) and maximum export rate (orange), export rates span a large range, mean baseline export half-life is 1.5 hours. **B:** Distribution of fold-change in amplitude over baseline. **C:** Barplot of fraction of genes selecting alternative models based BIC–full model (blue), model with cell cycle dependent export and constant splicing and transcription (purple), model with cell cycle dependent transcription and constant splicing and export (green), model with constant transcription, splicing, and export (red), for vast majority of the genes the full model is preferred. **D:** Polar histogram of the cell cycle phase at which a gene achieves the highest export rate. **E:** heatmaps of nuclear export during the cell cycle, the rates are normalized using max-min normalization, and the genes are arranged based on the timing of maximum export rate.

### Dynamics of chromatin accessibility during the cell cycle

The multiome profiling provides information about chromatin accessibility in addition to gene expression data for the same cells. This feature gave us the unique opportunity to investigate chromatin accessibility dynamics throughout the cell cycle by combining cell cycle phase estimation obtained using DeepCycle, and chromatin accessibility readout obtained by collecting all ATAC fragments mapped to a specific genomic region. We first examined the replicate correlation in the accessibility profiles for every gene (**Figure S6**) and filtered out those genes for which the correlation was weak (*ρ <* 0.75). Interestingly, we found that chromatin accessibility was highest for most genes in cells undergoing the S/G2 transition (**Figure 5A** and **5B**). This may be a consequence of DNA synthesis in combination with chromatin condensation during mitosis, *i*.*e*. genes that underwent DNA replication would contribute twice as much material, which serves as a readout for accessibility, followed by a global reduction of accessibility during mitosis. Furthermore, our data also suggests a preference in the order in which genes undergo DNA replication during the cell cycle (**Figure 5A**). To account for the ploidy, we normalized the accessibility readout by the total number of ATAC fragments detected in the nucleus. We found that the normalized accessibility profile revealed genespecific patterns of accessibility during the cell cycle. Averaging the normalized accessibility across all genes, we observed that there are two global peaks in relative accessibility, one during the G1/S and the second during the S/G2 transition (**Figure 5C** and **5D**). The normalized accessibility profile shows a prominent dip in global accessibility during mitosis, as expected due to chromatin condensation. When we examined the *z*-scored moving averages of unspliced, spliced, and normalized chromatin accessibility profiles for select genes that are known to achieve peak expression during the G2/M transition (Ccnb2, Nusap1, and Kif2c) we observed that their accessibility profiles vary significantly from one another (**Figure 5E**; **Figure S7**). This surprising observation indicates that the relationship between chromatin accessibility and transcription of individual genes might not be straightforward.

**Figure 5.**
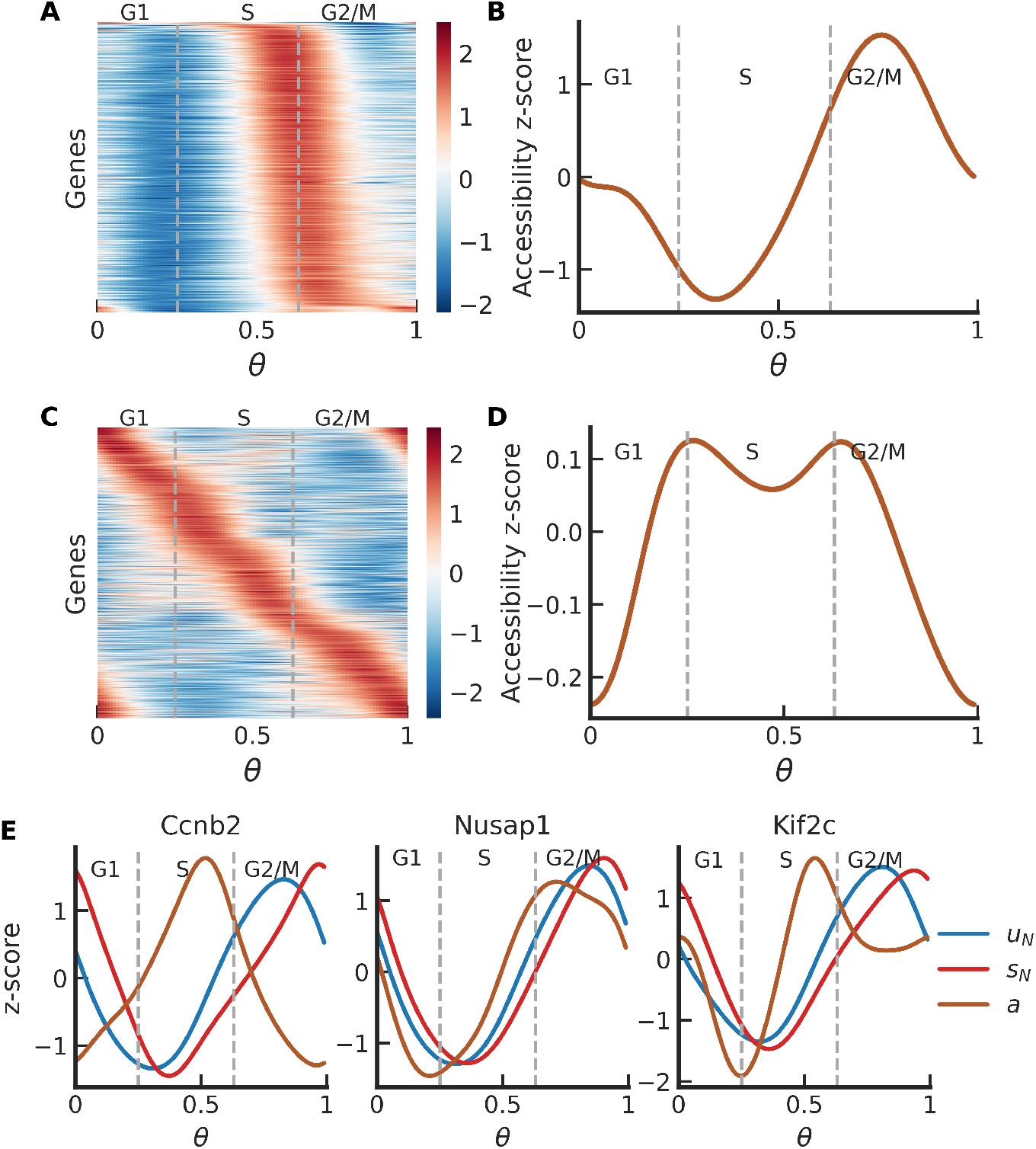
Chromatin accessibility during the cell cycle. **A:** Unnormalized chromatin accessibility against the cell cycle phase, *θ*, for genes with high replicate correlation, chromatin accessibility is maximum during the S/G2 transition which is consequence of the fact that there is twice as much DNA in these cells compared to that have not undergone DNA synthesis, the heatmap also suggests a preference in the order of which the genes undergo DNA replication. **B:** Average chromatin accessibility, across the genes show in panel A, is lowest at the beginning of S phase and highest at the beginning of G2 phase. **C:** Chromatin accessibility normalized for the total genetic material in cell, normalized accessibility of the genes is more evenly distributed during the cell cycle. **D:** Average accessibility is highest at the beginning of S and G2 phases and there is a global dip in accessibility during mitosis, likely due to chromatin compactification. **E:** Comparison of the dynamics of accessibility, spliced and unspliced profiles for select genes during the cell cycle; it is known that Ccnb2, Nusap1, and Kif2c expression levels peak in late G2/M phase, however, their accessibility peaks at different times during the cell cycle.

### Chromatin accessibility dynamics within TF footprints reveal key cell cycle regulators

ATAC-seq data has been instrumental in characterizing TF footprints within accessible chromatin regions [36]. Our single-nuclei multiome experiments, together with computational cell cycle sorting, allowed us to investigate the dynamic changes of TF footprints throughout the cell cycle. To that end, we first used a set of known Position Weight Matrices (PWMs) of 678 TFs to identify putative binding sites on the DNA within ATAC peaks found by CellRanger [37]. We then calculated TF footprints by combining all the observed Tn5 cuts at a given distance with respect to the predicted binding site (see Methods). This strategy produced average footprints for individual TFs and revealed distinct profiles, as illustrated for two representative factors Tfcp2l1 and Pml (**Figure 6A**). By repeating the same analysis considering nuclei in a specific cell cycle phase bin, we calculated how the footprints fluctuate with respect to the average throughout the cell cycle (**Figure 6B**). Interestingly, we observed an increase in chromatin accessibility around Tfcp2l1 binding sites during the G1-S transition, reaching a minimum during the G2-M transition (**Figures 6C** and **6D**). In contrast, the chromatin accessibility around binding sites of Pml exhibited a pronounced maximum during the G2-M transition (**Figures 6C** and **6D**). These changes in the TF footprints suggest potential variations in the binding activity of these TFs across different cell cycle phases. We then ranked all the investigated TFs based on the significance of their accessibility footprint changes (**Figure 6E**). Remarkably, among the top 40 TFs, several have been previously reported as regulators of the cell cycle (*e*.*g* : Chd1, E2f1, Nfya and Nrf1). This further supports the validity of our approach and highlights its potential to identify novel TFs involved in regulating transcription during the cell cycle. In summary, our method unveiled distinct cell cycle-dependent changes in TF footprints of chromatin accessibility, generating predictions of the specific phases when TFs are likely to actively interact with DNA and potentially regulate transcription during the cell cycle.

**Figure 6.**
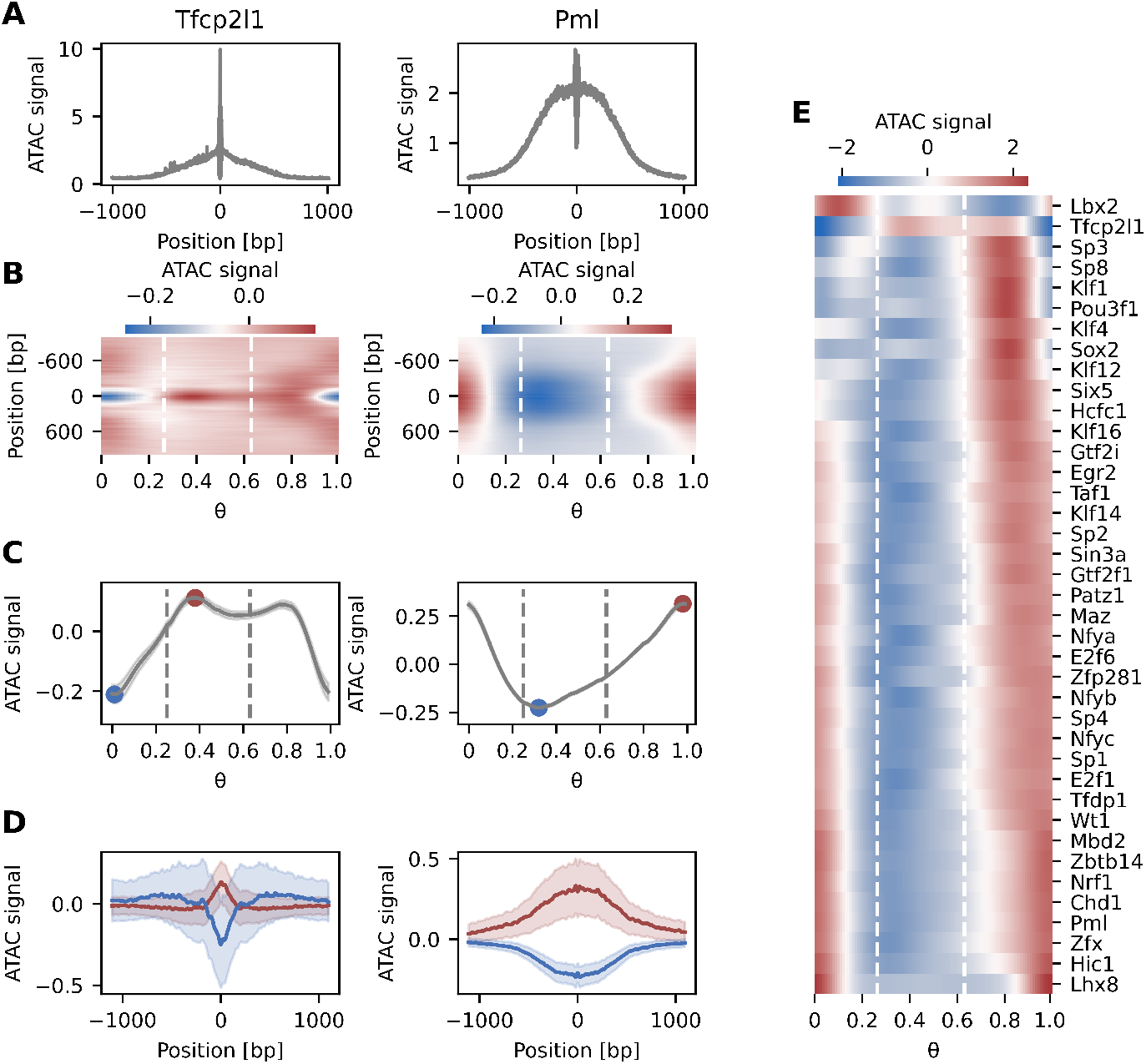
TF footprint dynamics during the cell cycle uncover key cell cycle regulators. **A:** Average ATAC-seq footprints of TFcp2l1 and Pml showing the normalized number of Tn5 cuts centered relative to predicted binding sites within ATAC peaks. **B:** Cell cycle dynamics of the Tfcp2l1 and Pml footprints as a function of the cell cycle phase *θ* relative to the average footprint depicted in A. White dashed lines delimit cell cycle phases in the following order: G1, S and G2/M. **C:** Changes in chromatin accessibility during the cell cycle averaged within a 100bp window centered at the TF binding site. Maximum and minimum accessibility are indicated in red and blue, respectively. Gray dashed lines delimit cell cycle phases in the following order: G1, S and G2/M. **D:** Relative TF footprint at the maximum (red) and minimum (blue) of accessibility as indicated in C. Shaded areas represent standard deviations. **E:** Changes in averaged chromatin accessibility around binding sites of the top 40 TFs for which the change is most significant. White dashed lines delimit cell cycle phases in the following order: G1, S and G2/M.

## Discussion

The RNA-velocity framework uses single-cell sequencing data (static information) to infer how gene expression changes over time and characterize gene expression dynamics in various biological contexts [14, 15]. DeepCycle uses the RNA-velocity framework to infer the cell cycle phase based on the unspliced and spliced levels of pre-selected cycling genes. Thus, DeepCycle reveals the oscillatory dynamics of gene expression at a very high temporal resolution [11]. In this work, using FourierCycle, we examined the relationship between the cell cycle and different processes involved in mRNA metabolism (transcription, splicing, mRNA degradation and mRNA nuclear export) and discovered oscillatory dynamics of mRNA metabolism, and waves of transcriptional and post-transcriptional regulation during the cell cycle.

To dissect different processes that affect the relationship between the cell cycle and mRNA metabolism, we constructed biophysical models that make genome-wide predictions for the cell cycle dependence of transcription, nuclear export, and degradation rates in mESCs. These models also give insights about the timing of up or down regulation of cellular biological processes during the cell cycle. Our predictions for the baseline rates, *i*.*e*. the cell cycle independent rates, were in agreement with the available literature [29, 30], validating our approach. It is relevant to point out that although the measurements for the rate parameters in the literature are derived from a variety of experiments, our biophysical models allow us to make predictions for all of these rate parameters based on a single experiment. Comparing the BIC score for alternate models we found that for most of the oscillating genes, the full model that incorporates cell cycle-dependent transcription, splicing and mRNA nuclear export is preferred. Importantly, we found that cell cycle-dependent changes in post-transcriptional regulation control the gene expression dynamics in proliferating mESCs to a greater extent than cell cycle-dependent transcription.

Our approach allowed us to examine the timing of peak rates, leading to the discovery of prominent waves of transcription, export, and degradation during the cell cycle. We found that nearly half of the oscillating genes achieved peak transcription during the early-G1 phase. This phenomenon has been previously shown in synchronized cell populations [9, 38, 39], to the best of our knowledge, this is the first time it is reported in unperturbed and unsynchronized mESCs. The peak transcription of the remainder of the genes is more evenly distributed throughout the cell cycle. The dynamics of nuclear export remains extremely poorly characterized, primarily due to lack of reliable transcriptomic measurements for the nucleus. Here, we debut a genomewide estimation of nuclear export rates and find that nearly 50% of the gene achieve maximal export in G2/M phase. Therefore, the disconnect between peak transcription rates and peak export rates hint at preferential nuclear retention of mRNA transcripts. We also observed that mRNA degradation waves are distinct from waves in transcription and export, with one wave occurring at around 25 mins of reentry into G1, another in late G1, and the third in early S phase. This results are in line with recent work by Krenning et al. that reported two distinct waves of mRNA degradation in RPE1 cells–an immediate wave within 20 mins after metaphase and a delayed wave roughly 80 mins after metaphase [40]. Taken together, our analysis reveals existence of distinct waves of transcription, mRNA nuclear export and mRNA degradation that occur at different points in the cell cycle to allow the cell to achieve tight temporal control over this complex process. Furthermore, to the best of our knowledge, this is the first time that genome-wide trasnscriptional and posttranscriptional waves have been reported during the cell cycle using scRNA-seq profiles. The advent of multiome single cell profiling has made it possible to simultaneously profile gene expression as well as chromatin accessibility at single cell resolution. This provided us with a unique opportunity to investigate accessibility dynamics during the cell cycle progression. One of the surprising findings that emerged from this work was the observation that genome-wide chromatin accessibility was the highest during the S/G2 transition, although one would expect that it would be highest during the S phase when the DNA is being duplicated. On the other hand, normalized accessibility does show a peak in accessibility during G1/S transition that stays relatively high during S phase, and undergoes a global dip during mitosis due to chromatin condensation. We were also able to probe gene-wise chromatin accessibilities and compare them with the gene expression readouts from snRNAseq data, as exemplified by Ccnb2, Nusap1, and Kif2c. We found hints of differential mechanisms involved in regulating the steps between chromatin accessibility and active transcription of associated genes.

Importantly, the paper describes an important extension of DeepCycle, as we validate that it can handle non only scRNA-seq data but snRNA-seq data as well. Our analysis of snRNA-seq enabled us to observe that the number of mRNA molecules in the nucleus increases during the cell cycle, which builds on previous observations that this also occurs in the cytoplasm [11]. This suggests that the local mRNA concentration stays constant during the cell cycle in both the cytoplasm and the nucleus. We also found extremely strong correlations in reads from scRNA-seq and snRNA-seq even though these are two completely different experimental protocols, giving us confidence in the quality of the measurements.

In conclusion, FourierCycle combines single-cell multiome sequencing and biophysical modeling to build a high-resolution map of the entire cell cycle transcriptome. This represents a powerful approach for genome-wide, single-cell studies of transcriptional dynamics throughout the cell cycle. Thus, FourierCycle and the underlining data we report here will serve as valuable resources for the community and enable dissection of regulatory mechanisms involved in oscillatory dynamics of mRNA metabolism during the cell cycle. Additionally, the scope of FourierCycle is not limited to in vitro models, as it could be easily extended to other systems, such as organoids or patient samples, and used to better understand cell cycle related mechanisms in development and disease biology.

## Limitations of the study

Single-cell sequencing technologies are inherently noisy, more so in the case of multiome sequencing where the genetic material from a single nucleus is shared between two experiments. We found extremely good concurrence in analysis between the two replicates snRNA-seq, which indicates high quality of our data. Recent work of Gorin et al. highlighted the shortcomings of RNA velocity framework, concerning both the underlying assumption of the biophysical model and preprocessing of the data. Our approach mitigates some of these limitations, specifically, we do not assume constant production and degradation rates, and our system does not have multiple cell identities, and most of the biological variabilitiy arises from cell cycle. DeepCycle is most effective for cells that are largely proliferative, and we would not recommend its use in cells that are quiescent, senescent, or with activate differentiation programs. Our future work will focus on overcoming some of these challenges and expanding the utility of DeepCycle further.

## Methods

### Cell culture

Mouse ES cells E14 were cultivated without feeders on 0.1% gelatine-coated dishes in DMEM high glucose (4.5 g/L) with GlutaMAX supplemented with 15% fetal calf serum heat inactivated, 0.001% B-mercaptoethanol, 1X non-essential amino acids, 1X NaPyr, 1X LIF, 1 mM MEK Inhibitor PD 0325901, and 3 mM GSK3 inhibitor CHIR99021.

### Nuclei isolation

Nuclei were isolated from 4X105 mouse ES cells using 100 *µ*L of cold lysis buffer (10 mM Tris-HCl (pH7.4), 10mM NaCl, 3 mM MgCl2, 0.1% Tween-20, 0.1% IGEPAL CA-630, 0.01% Digitonin, 1% BSA, 1mM DTT, 1 U/*µ*L RNase inhibitor). Cells were incubated for 5 min at 4°C, then resuspended in 1 mL chilled wash buffer (10 mM Tris-HCl (pH7.4), 10mM NaCl, 3 mM MgCl2, 0.1% Tween-20, 1% BSA, 1mM DTT, 1 U/*µ*L RNase inhibitor) and centrifuged at 500 rcf for 5 min at 4°C. Pellet was rinsed two more times with 1 mL wash buffer, for a total of 3 washes. The nuclei pellet was resuspended in 40 *µ*L Diluted Nuclei Buffer (1X Nuclei buffer (10X Genomics, PN-1000280), 1 mM DTT, 1 U/*µ*L RNase inhibitor), and passed through a 40 *µ*m Flowmi Cell Strainer (Merck).

### Single nuclei library preparation and sequencing

Library preparation was performed at the GenomEast platform at the Institut de Génétique et de Biologie Moléculaire et Cellulaire. Transposition was performed using the 10X Genomics kit (PN-1000280) and nuclei were loaded into the 10X Genomics Chromium Controller with a target capture of 6,000 single nuclei. Libraries for the ATAC and 3′ mRNA were produced according to the Chromium Next GEM instructions (10X Genomics, Document CG000338 Rev F), and sequenced using the Illumina NextSeq 2000. Libraries were sequenced on an Illumina NextSeq 2000 sequencer as paired-end 28 + 85 base reads for 3′ mRNA, 50 base reads for ATAC. Image analysis and base calling were performed using RTA version 2.7.7 and BCL Convert version 3.8.4.

### Data Processing

Raw data was processed using CellRanger (version 7.0.1), cellranger count option with the 10X reference genome mm10-2020-A, for demultiplexing, barcoded processing, and gene counting. Next we used velocyto (version 0.17.17, python implementation), run10x function along with the 10X reference genome to map the reads to intronic and exonic regions to obtain the unspliced and spliced quantification. We then used scVelo (version 0.2.3) for smoothing the unspliced and spiced quantifications with these hyperparameters: `min shared counts=20`, `n pcs=30`, `n neighbors=30`. Lastly We ran DeepCycle on the processed data to infer the cell cycle phases.

### Biophysical model

We described the dynamics of the unspliced and the spliced mRNA for every gene with respect to the transcriptional phase during the cell cycle using a system of coupled ordinary differential equations,

Cell:

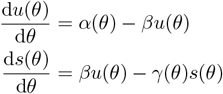

Nucleus:

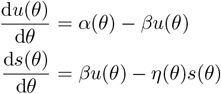

where *α* denotes the synthesis rate, *β* denotes the processing or the splicing rate, *η* denotes the nuclear export rate, and *γ* denotes the degradation rate. For every gene, we allow the synthesis, export, and degradation rates to explicitly depend on the cell cycle phase, *θ*. Since the dynamics in question is cyclic in nature, *u, s, α, η*, and *γ* will be periodic functions of *θ*. We used the Fourier approximation to solve this system of ordinary differential equations (details in supporting information).

### Model fitting

The optimization was performed using the `optimize.minimize` package from `scipy` module in python. We used loss function to be minimized was,

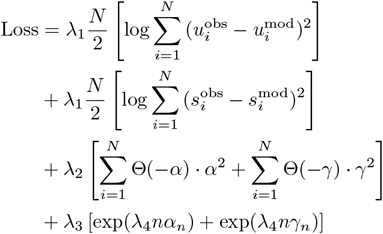

where *u*_*i*_ and *s*_*i*_ denote the observed unspliced and spliced reads, whereas 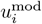 and 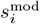 denote the model predictions of the unspliced and spliced reads in the *i*^th^ cell.

For every gene we calculated the explained variance,

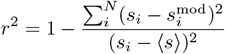

where ⟨*s*⟩ denotes the average spliced molecules across all the cells. We set a threshold of 20% and excluded all genes below this value.

### Chromatin accessibility analysis

To compute chromatin accessibility dynamic profiles for each gene, we used the 10x gene annotation (Ensembl98) and extracted genomic regions from 2000 bp upstream of the TSS to the end of the gene. Then, The 10X “iscell” barcode fragments were extracted and intersected with the above genomic regions with bedtools coverage tool (v2.30.0). This resulted in two matrices, one for each replicate, containing the number of ATAC fragments mapped to each gene in every cell. Sorting cells according to their cell cycle phase, *θ*, and smoothing the number of ATAC fragments across cells produced chromatin accessibility profiles for each gene during the cell cycle. Only genes showing high replicate correlation (*ρ >* 0.75) between the chromatin accessibility profiles in each replicate were retained.

To determine TF footprints within accessible chromatin, we initially identified putative TF binding sites in ATAC peaks. To achieve this, we run MoteEvo [41] with a set of known Position Weight Matrices (PWMs) of 678 TFs obtained from the SwissRegulon database [42]. We then calculated TF footprints for each cell by aggregating all observed Tn5 cuts at a given distance from the predicted binding site within a window of ± 1 Kbp. Cell cycle-dependent TF footprints were computed by smoothing single-cell data sorted according to their cell cycle phase, *θ*. Additional smoothing along the relative genomic coordinates with respect to the TF binding sites was performed. Finally, relative TF footprints were calculated by subtracting the average TF footprint across the cell cycle.

## Resources

Processed data:

https://doi.org/10.5281/zenodo.10462840

Github:

https://github.com/MolinaLab-IGBMC/

FourierCycle

Web server:

https://molina.igbmc.science/

FourierCycle-Home.html

## Acknowledgments

We are grateful to Manuel Mendoza, Bertrand Séraphin, Georg Stoecklin, and Jingkui Wang for their thorough review and constructive criticism, which greatly improved the overall quality of this manuscript. This work of the Interdisciplinary Thematic Institute IMCBio+, as part of the ITI 2021-2028 program of the University of Strasbourg, CNRS and Inserm, was supported by IdEx Unistra (ANR-10-IDEX-0002), and by SFRI-STRAT’US project (ANR-20-SFRI-0012) and EUR IM-

CBio (ANR-17-EURE-0023) under the framework of the France 2030 Program. This work was also supported by a fellowship from L’Institut d’Études Avancées de l’Université de Strasbourg (USIAS). Library preparation and sequencing were performed by the GenomEast platform, a member of the France Génomique consortium (ANR-10-INBS-0009). Icons used in Figure 1 made by dDara (cell), Dimitry Mirolibov (DNA), Mayor Icons (anaphase), and Freeplk (cytokinesis) from www.flaticon.com.

## Author contributions

MKN and NM conceptualized the study, DS and MC performed the experiments, MKN, AR, TY, and NM performed bioinformatics analyses, MKN and AR developed the code for biophysical modeling, OT developed the web server, MKN, DS, and NM wrote the manuscript, and NM led the research, secured funding, and provided supervision.

## Supplementary figures

**Figure S1.**
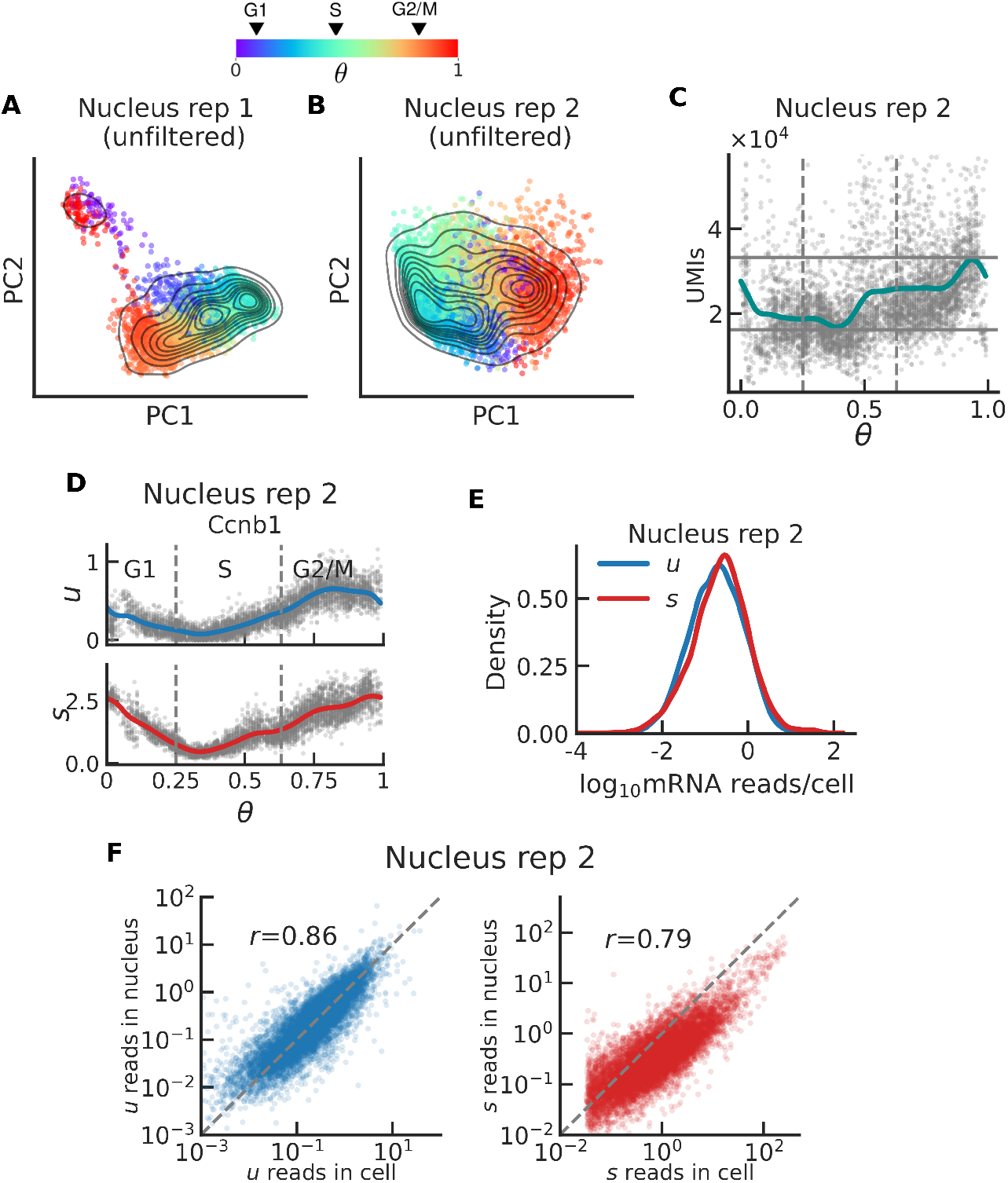
Data characteristics of single-nucleus sequencing data. **A** and **B:** Principal components analysis of unfiltered snRNA-seq data. **C:** UMIs as a proxy for the total mRNA content in the nucleus (replicate 2), against the cell cycle phase *θ*. **D:** Expression dynamics of the unspliced and spliced reads for Nusap1 in the nucleus (replicate 2). **E:** Distribution of unspliced (blue) and spliced (red) reads per cell, across all genes in the nucleus (replicate 2). **F:** Correlation between the reads in the cell versus those in the nucleus (replicate 2) for the unspliced (blue) and spliced (red) molecules respectively.

**Figure S2.**
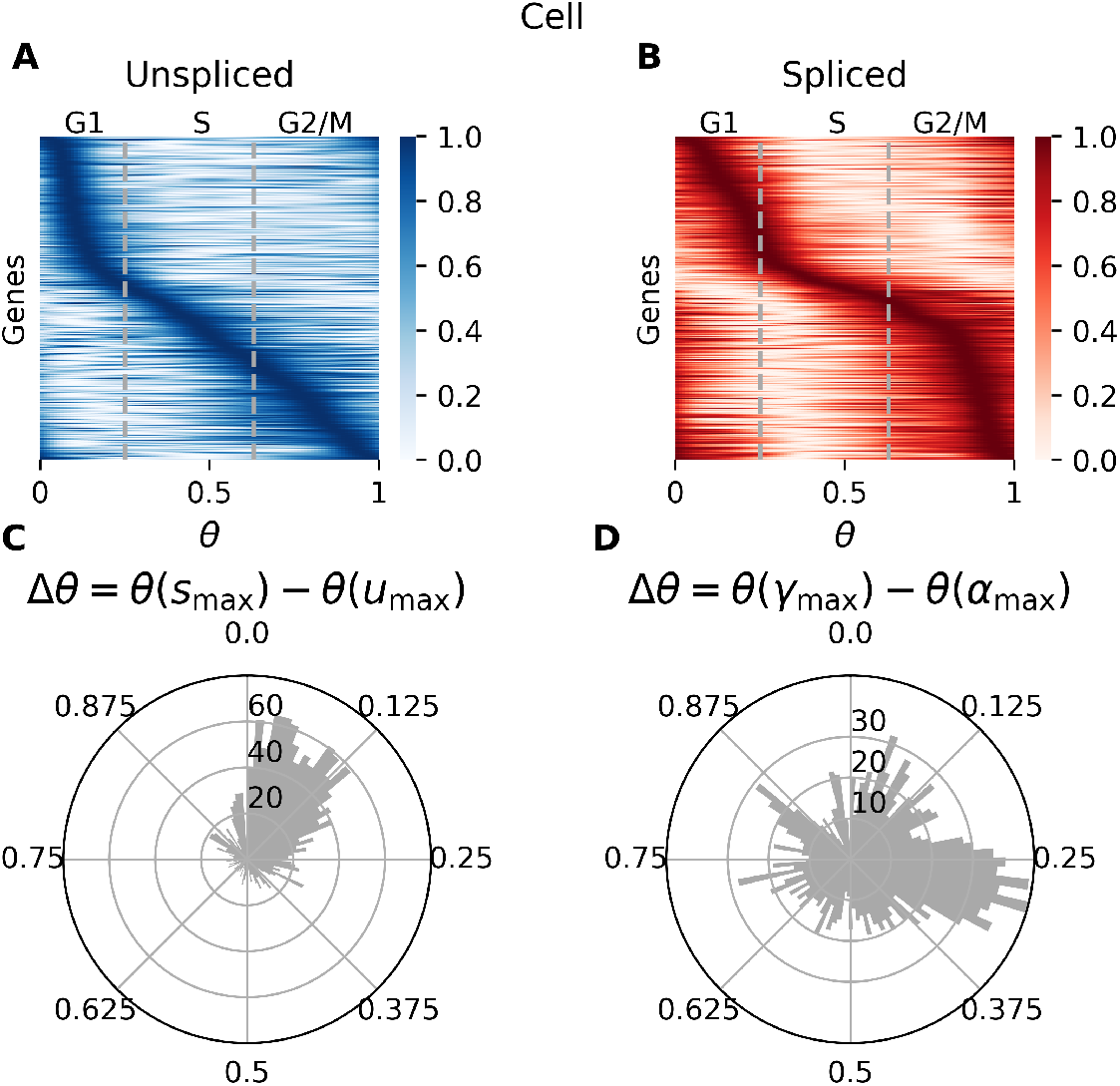
Heatmaps of gene expression and histograms of time differences for the cell. **A** and **B:** Heatmaps of normalized, using max-min normalization, of unspliced and spliced quantifications of the genes filtered based on the explained variance cutoff. **C:** Polar histograms of time difference between maximum unspliced and maximum spliced **D:** Polar histograms of time difference between maximum transcription and maximum degradation for scRNA-seq data.

**Figure S3.**
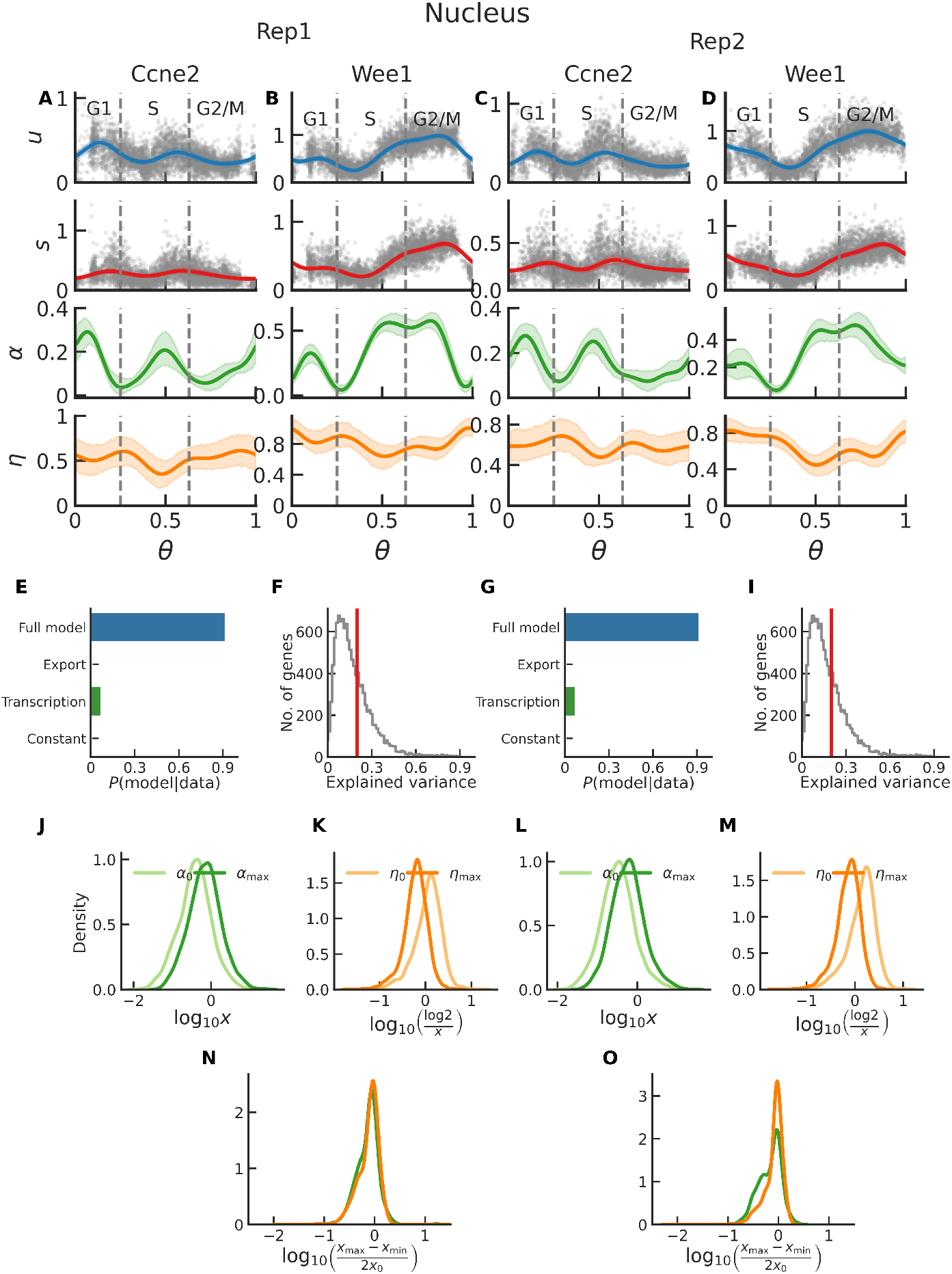
Kinetic modeling results for single-nucleus sequencing data. **A-D:** Number of unspliced reads (*u*), number of spliced reads (*s*), transcription rate (*α*), export rate (*η*) as function of the cell cycle phase *θ* for representative genes Ccne2 and Wee1 (multiome snRNA-seq data replicates 1 and 2). **E** and **G:** Barplot of fraction of genes selecting alternative models based on the Bayesian Information Criterion for replicates 1 and 2 respectively. **F** and **H:** Histogram of the explained variance in spliced counts across all the genes for replicate 1 and 2 respectively; red line is at *x* = 0.2, the threshold set for filtering genes. **J** and **L** Distribution of transcription rate, in arbitrary units, *α*_0_ denotes baseline transcription (light green) whereas *α*_max_ denotes maximum transcription rate (green) for replicates 1 and 2 respectively. **K** and **M:** Distribution of export half lives, in hours, across filtered genes, *η*_0_ denotes baseline export rate (light orange) whereas *η*_max_ denotes maximum export rate (orange) for replicates 1 and 2 respectively.

**Figure S4.**
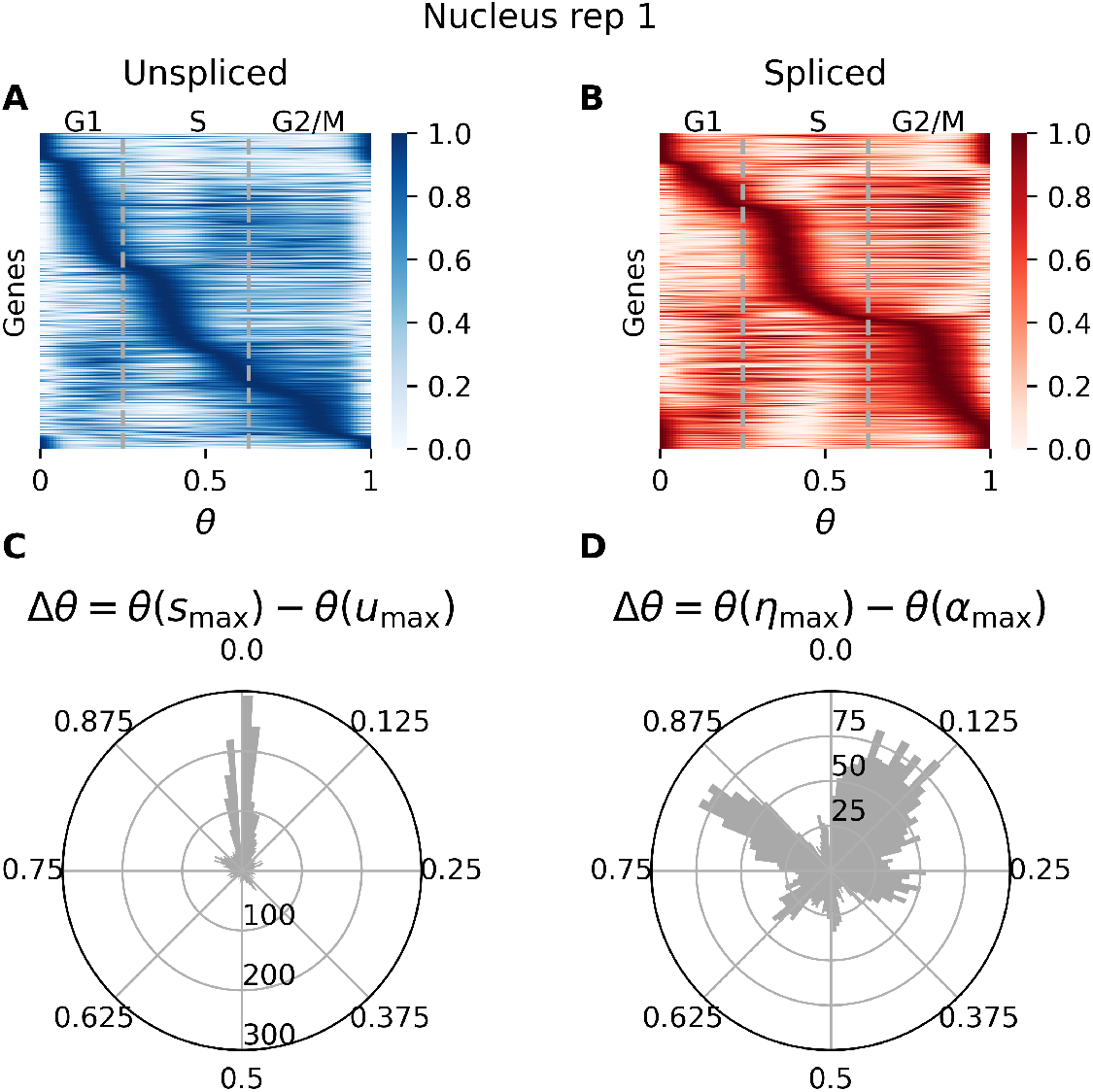
Heatmaps of gene expression and histograms of time differences for the cell. **A** and **B:** Heatmaps of normalized, using max-min normalization, unspliced and spliced of the genes filtered based on the explained variance cutoff. **C:** Polar histograms of time difference between maximum unspliced and maximum spliced. **D:** Polar histograms of time difference between maximum transcription and maximum export for snRNA-seq data.

**Figure S5.**
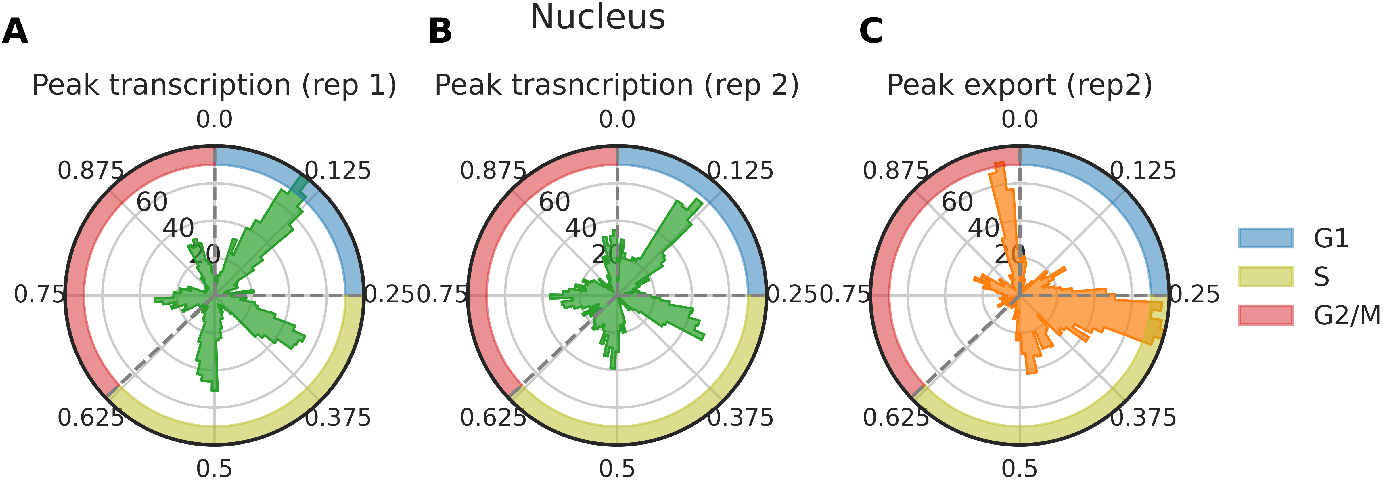
Waves of transcription and export in the nucleus. **A** and **B:** Polar histograms for timing of peak transcription for snRNAseq data (replicates 1 and 2) and **C:** Polar histograms of timing of peak export (replicate 2, replicate 1 is in Figure 5D)

**Figure S6.**
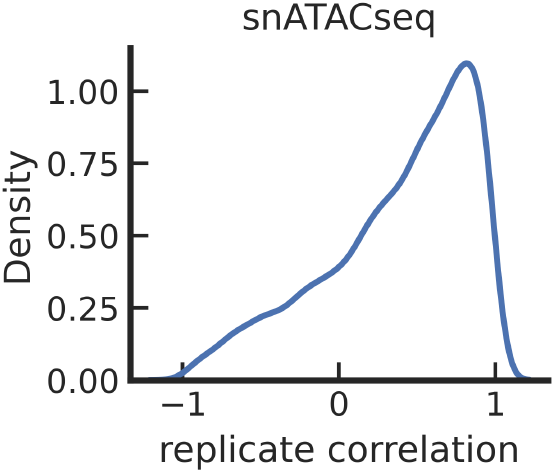
Replicate correlation for snATACseq data.

**Figure S6.**
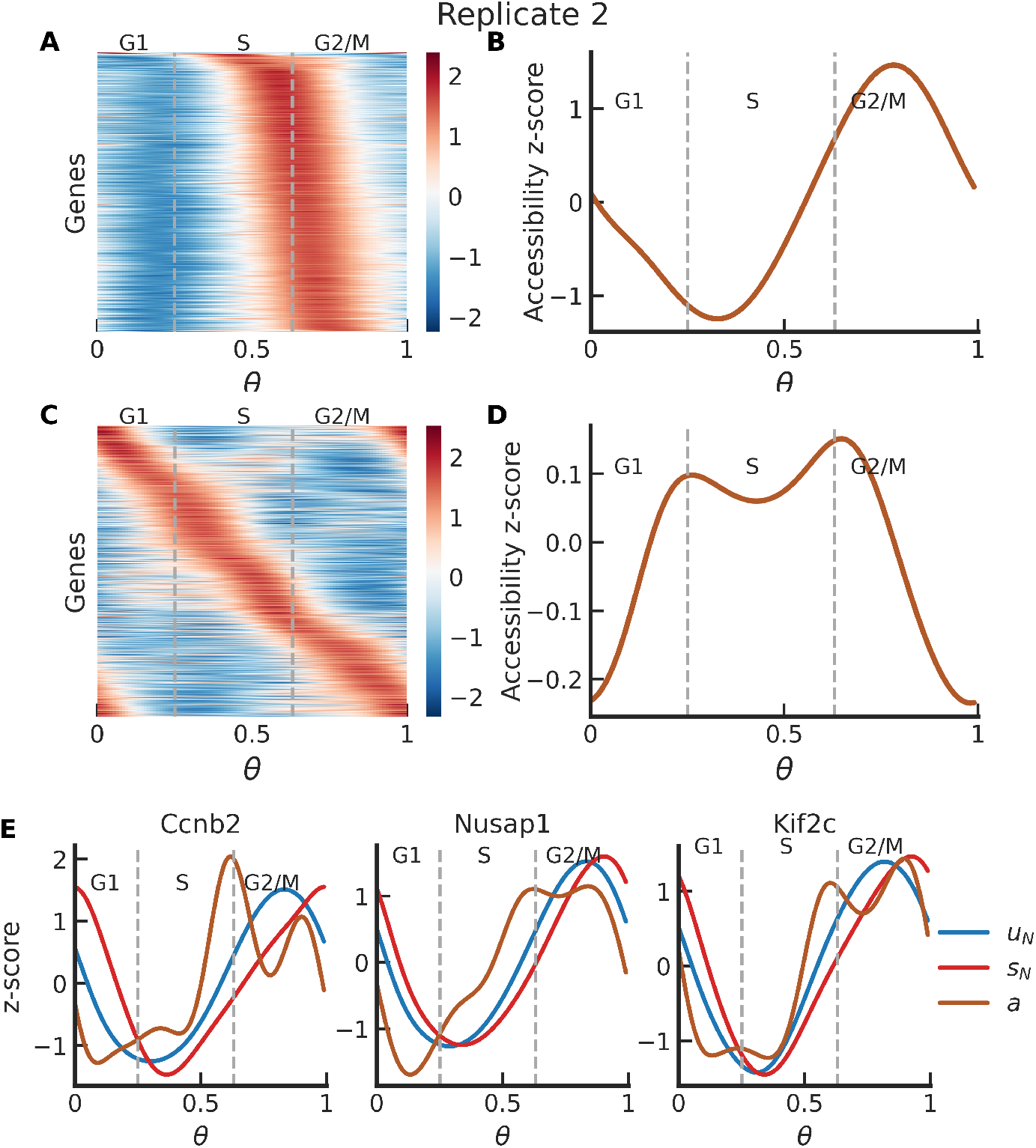
Chromatin accessibility during the cell cycle. Same as panels in figure 6 for replicate 2.

### Supporting information

#### Biophysical model

We model the dynamics of mRNA metabolism by expressing the steps involved—transcription, splicing, degradation—as a system of coupled ordinary differential equations.

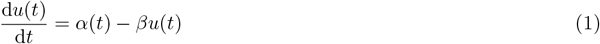

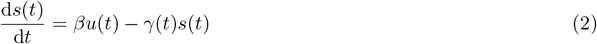

*α*: synthesis rate

*β*: splicing rate

*γ*: degradation rate

#### Unspliced solution

We assume that *u*(*t*) and *α*(*t*) are periodic functions. Thus, we can express *u*(*t*) and *α*(*t*) as sums of Fourier series.

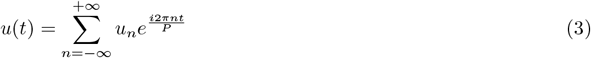

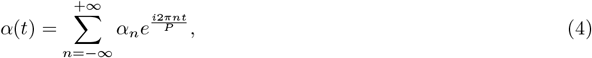

wehre *P* : the period of oscillation. Let 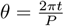 such that θ ∈ (0, 1)

Thus, we have

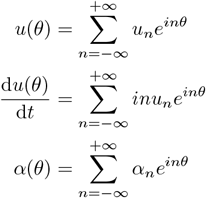

Using these in equation (1), we get,

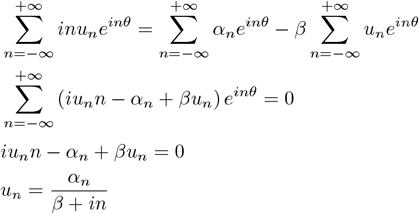

The Fourier approximation for *u*(*θ*)

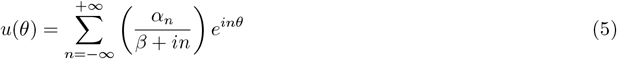

When performing numerical optimization, it is convenient to have real-valued parameters, *i*.*e*. Fourier coefficients for *α*(*θ*) and *γ*(*θ*). Fourier series for *α* with real-values co-efficients can be written as

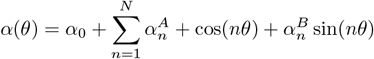

We can recover the complex co-efficients

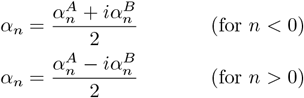

#### Spliced solution

Akin to *u*(*θ*) and *α*(*θ*), we expect *s*(*θ*) and *γ*(*θ*) also to be periodic functions. The Fourier series expansion of *s* and *γ* (as a function of *θ*) is,

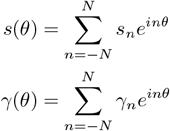

Using these expressions in (2),

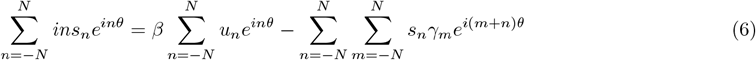

In the last term in the above equation, let *k* = *m* + *n*

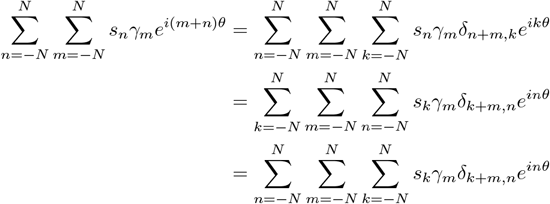

Using this in (6)

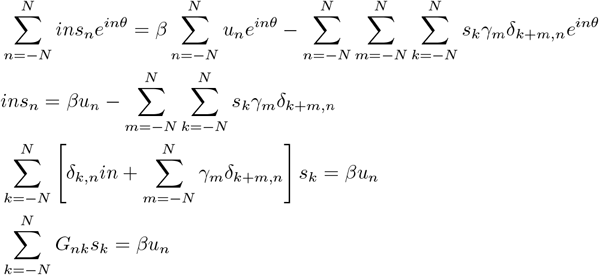

We can express this as a matrix equation

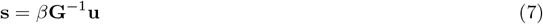

where **G** is a (2*N* + 1) × (2*N* + 1) matrix:

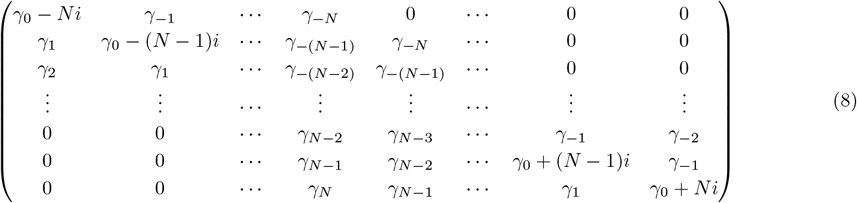

#### Model fitting

The optimization was performed using `scipy.optimize.minimize`, we used the following loss function for the optmization,

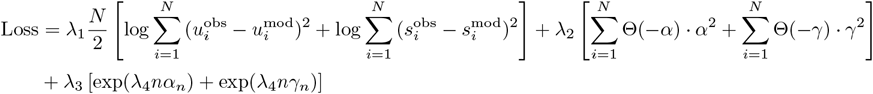

